# Distinct Differences in Gene Expression Profiles in Early and Late Stage Rhodesiense HAT Individuals in Malawi

**DOI:** 10.1101/2022.11.28.518140

**Authors:** Peter Nambala, Julius Mulindwa, Harry Noyes, Joyce Namulondo, Oscar Nyangiri, Enock Matovu, Annette MacLeod, Janelisa Musaya, TrypanoGEN+ Research Group as Members of the H3Africa Consortium

## Abstract

*T. b. rhodesiense* is the causative agent of rhodesian Human African trypanosomiasis (r-HAT) in Malawi. Clinical presentation of r-HAT in Malawi varies between the different foci and differs from East African HAT clinical phenotypes. The purpose of this study was to gain more insights into the transcriptomic profiles of patients with early stage 1 and late stage 2 HAT disease in Malawi. Whole blood from individuals infected with *T. b. rhodesiense* was used for RNA-Seq. Control samples were from healthy trypanosome negative individuals matched on sex, age range, and disease focus. Illumina sequence FASTQ reads were aligned to the GRCh38 release 84 human genome sequence using HiSat2 and differential analysis was done in R using the DESeq2 package. XGR, ExpressAnalyst and InnateDB algorithms were used for functional annotation and gene enrichment analysis of significant differentially expressed genes. RNA-seq was done on 25 healthy controls and 23 r-HAT case samples of which 3 case samples were excluded for downstream analysis as outliers. 4519 genes were significantly differentially expressed (p adjusted <0.05) in individuals with early stage 1 r-HAT disease (n = 12) and 1824 genes in individuals with late stage 2 r-HAT disease (n = 8). Enrichment of innate immune response genes through neutrophil activation was identified in individuals with both early and late stages of the disease. Additionally, lipid metabolism genes were enriched in late stage 2 disease. We further identified uniquely upregulated genes (log2 Fold Change 1.4 - 2.0) in stage 1 (ZNF354C) and stage 2 (TCN1 and MAGI3) blood. Our data brings new insight into the human transcriptome landscape during *T. b. rhodesiense* infection. We have further identified key biological pathways and transcripts during stage 1 and stage 2 r-HAT. Lastly, we have identified potential diagnostic biomarkers that may be used for staging of r-HAT disease.

## Introduction

Human African Trypanosomiasis (HAT) is a protozoan disease endemic in sub-Saharan Africa and caused by *Trypanosoma brucei gambiense* (Tbg) and *Trypanosoma brucei rhodesiense* (Tbr). Tbg causes chronic HAT (g-HAT) or sleeping sickness in West and Central Africa with domesticated animals sometimes acting as intermediate hosts (1). Whereas, Tbr causes an acute HAT (r-HAT) disease phenotype and is endemic in Southern and Eastern Africa where wild life and domesticated animals are the major intermediate hosts for disease transmission (2). Tsetse flies of the genus *Glossina* are the vectors of HAT transmission (3). Human Tbr infections are characterized by a hemolymphatic stage 1 (early) and meningoencephalitic stage 2 (late) disease. Parasite invasion of central nervous system is a classic characteristic of stage 2 r-HAT disease and if untreated, patients die due to a dysfunctional immune response in the central nervous system (4). Early diagnosis of r-HAT is key in reducing disease burden and mortality (5). Unfortunately, diagnosis of r-HAT in endemic areas is dependent on insensitive microscopic examination of blood and the invasive lumber puncture for collection of cerebral spinal fluid (CSF) used in r-HAT staging (6, 7).

There are variations in the clinical presentation of r-HAT that have been observed in endemic countries (8). For instance, r-HAT cases in Uganda frequently present with a more acute clinical presentation compared to r-HAT cases in Malawi that tend to present with a chronic disease phenotype (9). In Malawi HAT cases are reported at the interface of wildlife reserves with human settlements in Nkhotakota and Rumphi (10). Most cases in Nkhotakota present with a chronic stage 1 disease whereas in Rumphi most cases present with acute stage 2 disease (10, 11). Variations in clinical presentation of r-HAT have also been reported between Malawi and Uganda cases and this is associated with parasite genetic diversity and human inflammatory cytokine response (8, 12).

Previously, human transcriptome analysis of blood from r-HAT patients in Uganda identified functional enrichment of genes involved in innate immune response pathway to be the most differentially expressed (13). These genes include interleukin 21 *(IL21)*, interleukin 1 receptor *(IL1R)*, and tumour necrosis factor alpha *TNFA*, as well as immunoglobulin heavy chain variable and classical complement pathway genes, (13). Whereas, upregulated transcripts in the CSF of stage 2 HAT patients were predominantly coding for genes involved in neuro activation and anti-inflammatory, the study identified *IGHD3–10, C1QC* and *MARCO* genes as having a fivefold change in stage 1 r-HAT cases compared to healthy controls (13). The dual (host and parasite) transcriptome analysis of transcriptomes found in Ugandan r-HAT samples are unlikely similar to other r-HAT endemic counties such as Malawi. Firstly, Tbr parasite isolates in East Africa are genetically different from Malawi isolates (8). Secondly, clinical presentation of r-HAT in Malawi is more chronic than the acute disease observed in East Africa (9). Thirdly, there is a high level of human genetic diversity between Uganda and Malawi, which might affect human response to diseases (14). Lastly, association studies have found a protective effect of *APOL1* gene polymorphisms in r-HAT disease outcome in Malawi population (15), in contrast to mixed results from association studies of *APOL1* with r-HAT disease outcome in Ugandan population (16, 17). In this study, we examined the differences in the human blood gene expression profiles of r-HAT patients in Malawi. Our results add to the current understanding of the human response to r-HAT disease and have led to identification of potential blood markers for staging of r-HAT.

## Results

### RNA-Seq Sample attributes

Samples were collected at Rumphi and Nkhotakota district hospitals during a HAT surveillance as we had previously described (10). In Rumphi district, a total of 37 r-HAT positive cases and 25 corresponding r-HAT negative controls were recruited **(Table 1)**. Of the 37 r-HAT positive individuals, 26 (70.3%) were males and 11 (29.7%) were females. The mean age of the cases and controls were 34.9±17.2 years and 36.0±17.7 years respectively.

**Table 1.**
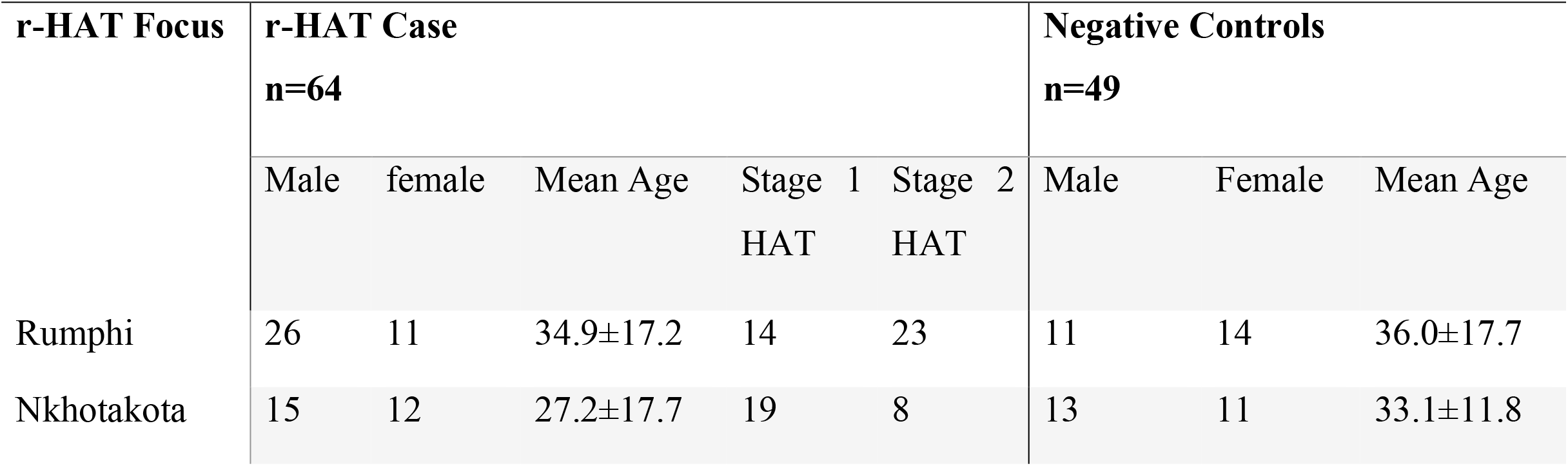
Demographic details of recruited study participants

In Nkhotakota district, 27 r-HAT cases were recruited and together with 24 corresponding negative controls **(Table 1)**. Among the cases, 15 (55.6%) were males and 12 (44.4%) females. The mean age of the cases and controls were 27.2±17.7 years and 33.1±11.8 years respectively. The HAT status of the participants was confirmed by microscopic examination of thick blood smears at recruitment sites and by PCR to detect the *SRA* gene of Tbr parasites as previously described (10).

RNA-Seq was done on 23 r-HAT cases and 25 healthy control blood samples with RNA concentration >1µg **(Table S1)**.

### Distinct differences in r-HAT cases and control transcriptome profiles

To examine for differences between the blood transcriptomes of individuals infected with Tbr parasites compared with healthy controls, we performed a principal component analysis (PCA) in DEseq2 (18). Three cases were identified as outliers by PCA and removed from downstream analyses. The results show that transcriptomes in individuals infected with Tbr parasites were clearly distinguished from healthy controls on a plot of principal components 1 and 2 (**Fig 1A**). We also observed a stratification when simultaneous comparison of female and male r-HAT cases with corresponding health controls was made using Euclidean distance correlation (**Fig S1A**). Lastly, we observed significant differentially expressed genes (DEGs) between stage 1 and stage 2 samples against controls **(Fig 1B and Fig S1B)**. Since clinical presentation or r-HAT in Malawi is focus dependent (10), we next compared transcriptome of infected individuals in Nkhotakota focus against infected individuals in Rumphi focus. No genes were significantly differentially expressed in infected individuals between the two r-HAT foci.

**Fig 1.**
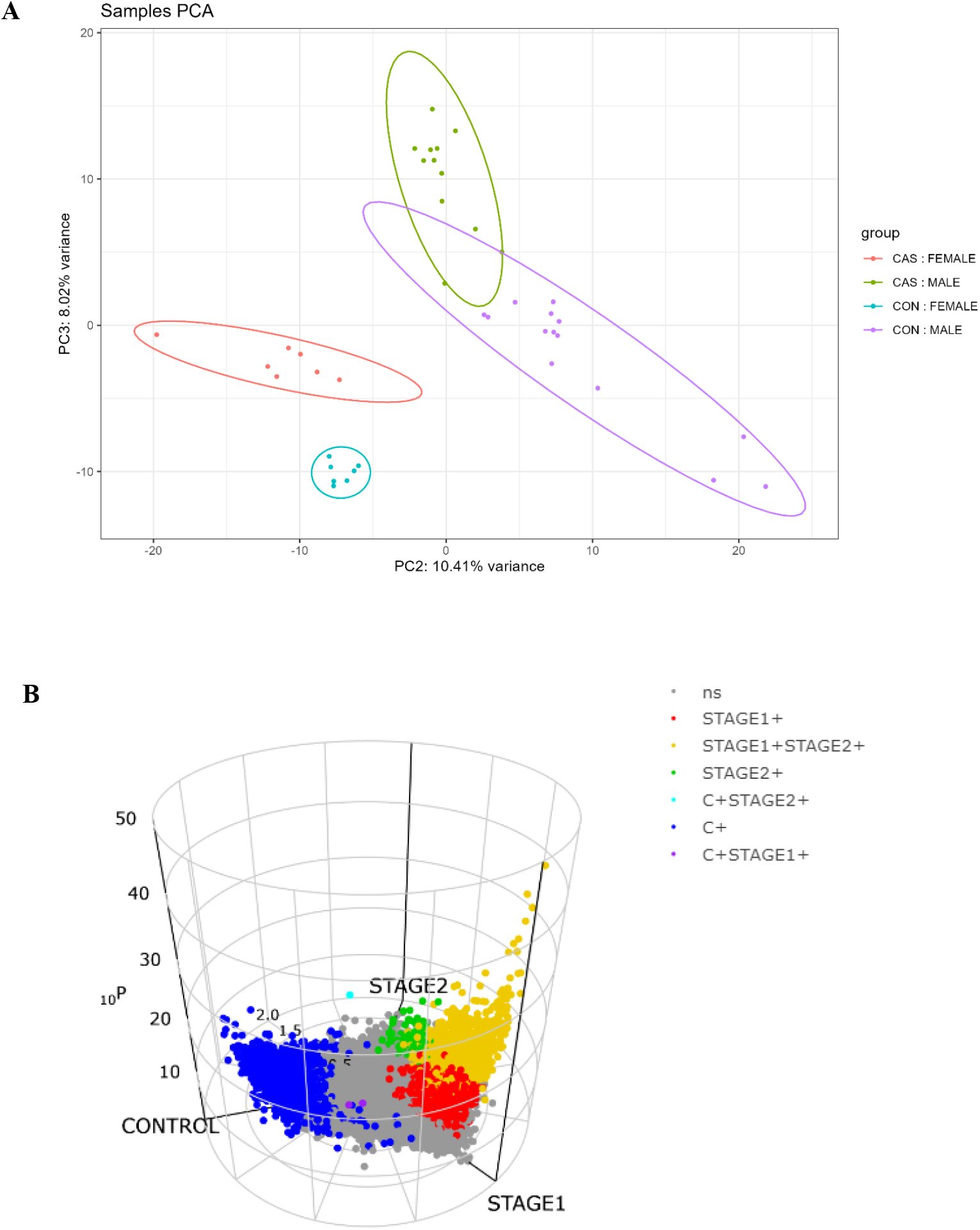
Stratification of Differentially Expressed genes (DEGs).**(A)** Principal component analysis (PCA) values for r-HAT cases vs healthy controls grouped into males and females. **(B)** 3D volcano plot showing distribution and relationship of DEGs in Stage 1, Stage 2 and Controls. Grey dots represent non-significant genes, dark blue dots are genes expressed in controls only, red dots are genes expressed in stage 1 only, green dots are genes expressed in stage 2 only, orange are genes expressed in both stage 1and stage 2, purple dots are genes expressed in controls plus stage 1 and light blue dots are genes expressed in controls plus stage 2.

### Innate Immune Response Transcripts are Elevated in Stage 1 Patients

Given the differences observed in the number of DEGs between HAT stage 1 and stage 2 blood relative to controls, we next sought to identify those genes that are significantly enriched in individuals with stage 1 r-HAT disease. First, differential transcriptome analysis was done in stage 1 cases against healthy controls using DeSeq2. A total of 4519/47546 (9.50%) genes were significant differentially expressed between stage 1 cases and healthy controls with adjusted p<0.05 (padj<0.05) (**Fig 2A**). Of the 4519 genes, 54.3% (2454/4519) coded for proteins, 32.2% (1457/4519) for lncRNA and 13.5% (608/4519) for various gene types which include miRNA, snRNA, snoRNA, scaRNA, miscRNA, Tyrosine protein kinase (TEC), immunoglobulins, T cell receptor and Mt-RNA (**Fig S2A**). Of the 2454 protein coding genes, 64.6% (1585/2454) were upregulated (log2 fold change, log2FC > 1), 8.2% (201/2454) were down regulated (Log2FC < -1) and 27.2% (668/2454) were neither upregulated nor downregulated compared to healthy controls. Among upregulated genes: *BMP6, ENOSF1, EXOSC9, SMARCB1, LCORL, EMC9, C12orf73, IFI16* were significantly expressed with padj<10e-15 and log2FC > 2. BMP6 plays a critical role in cell proliferation and type II cytokine regulation through JAK2 signalling pathway (19). IFI16 has a critical role in the interaction between the innate immune system and cellular transcriptional regulation through pattern recognition of pathogens. Additionally, upregulation of immunoglobulin light chains (IGKs, IGLs) and immunoglobulin heavy chains (log2FC 2.0 – 6.0) were identified. IGKs and IGLs are involved in activation of mast cells and neutrophils which results in the release of various pro-inflammatory mediators (20); whereas immunoglobulin heavy chains are central in presentation of antigens (21). All T cell receptor transcripts were downregulated (log2FC -1.0 to -2.1) in Stage 1 cases compared to healthy controls.

**Fig 2.**
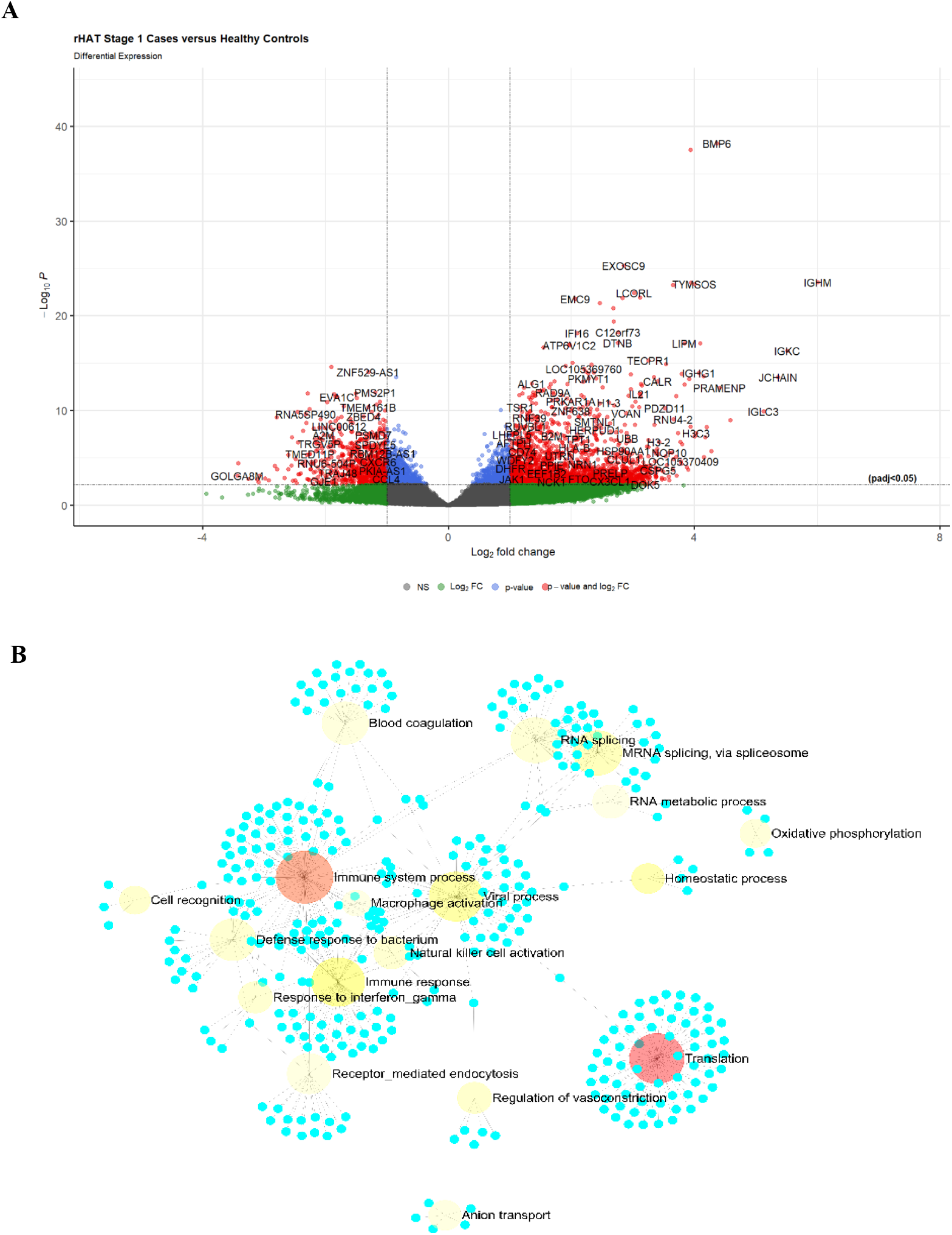
DEGs and network analysis in Stage 1 case. **(A)** Volcano plot showing genes that were significant (padj< 0.05) DEG, upregulated (log2FC > 1.0) and downregulated (log2FC < -1). **(B)** ExpressAnalyst network graph of upregulated protein coding genes. The root of the nodes was color coded according to significance with light yellow representing less significant and red more significant. Translation and immune system process were the most enriched biological pathways in stage 2 blood.

*T*.*b. rhodesiense* infections are known to disrupt the circadian rhythm which results in sleep disturbance in infected individuals (4, 22-24). For this, we observed that circadian rhythm *CIPC* (*clock interacting pacemaker*) was differentially expressed (padj<1.59E-6) and down regulated (log2FC -1.9) suggesting a disruption of circadian rhythm in patients with stage 1 disease compared to healthy individuals. However, *period circadian regulator 1* (*PER1*) transcripts were not differentially expressed but were upregulated (log2FC 1.7). *clock circadian regulator* (*CLOCK*) transcripts which is also central in circadian rhythm was neither differentially expressed nor upregulated.

Functional annotation of the principal component gene ontology (25), identified immune system function as having the most enriched genes with high loadings on the selected principal components **(Table S2)**. Immune effector process, neutrophil activation, neutrophil degranulation, neutrophil activation involved in immune response and neutrophil mediated immunity were among differentially expressed immunological functions (p-value<10E-12). Neutrophils have been previously shown to have a fundamental role in innate immune response against trypanosome parasites (26, 27).

To determine other biological processes enriched during stage 1 of Tbr infection, upregulated genes were analysed in ExpressAnalyst using the PANTHER biological process database (28, 29). This identified 18 biological process which include immune system process, immune response, macrophage activation, natural killer cell activation, response to interferon gamma, cell recognition, receptor mediated endocytosis and blood coagulation **(Fig 2B)**. The blood coagulation system may be activated by pro-inflammatory cytokines and modulate inflammatory response to blood pathogens (30).

### Enrichment of Lipid Metabolic Process Pathway in stage 2 r-HAT Cases

To determine blood transcriptomes that were enriched in stage 2 patients, we compared stage 2 blood against heathy controls. There were 1824/37922 (4.81%) significant DEGs (padj<0.05) of which 850/1824 (46.6 %) coded for proteins, 643/1824 (35.3 %) for lncRNA and 331/1824 (18.1 %) for various gene types **(Fig 3A and Fig S1B)**. Additionally, 75/850 (8.8%) of the protein coding genes were highly upregulated (log2FC 2.0 - 4.3) relative to healthy controls with *BMP6, ENOSF1, IFI16, SMARCB1* and C12orf73 genes significantly expressed with padj<9.79E-10. Whereas 17/850 (2.0%) protein coding genes were highly downregulated (log2FC -2.0 to -3.7) with *UGT2B28* significantly expressed (padj<9.81E-5). All upregulated gene (375/850) were analysed in ExpressAnalyst to identify pathways enriched in the biological process in PANTHER biological process database. This identified translation (padj<9.19E-6), immune system process (padj<3.59E-4) and immune response (padj<0.004) as the most significant enriched biological pathways in stage 2 r-HAT cases **(Fig 3B)**.

**Fig 3.**
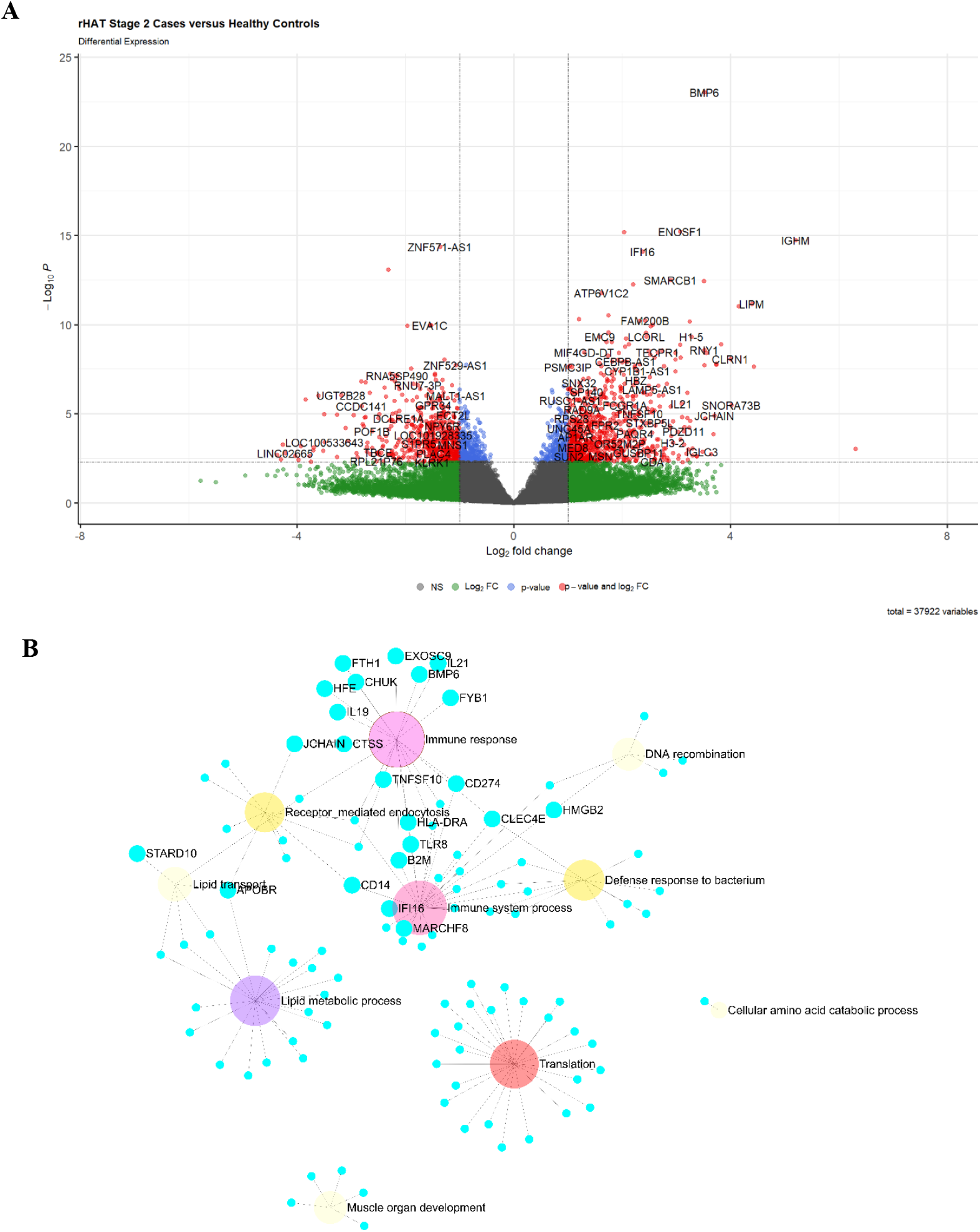
DEGs and network analysis in Stage 2 case versus healthy controls. **(A)** Volcano plot showing **s**ignificant DEGs (padj< 0.05) that were upregulated (log2FC > 1.0) and downregulated (log2FC < -1.0). **(B)** ExpressAnalyst network graph of protein coding genes that were upregulated in stage 2 blood relative to controls. The root of the nodes was color coded according to significance with light yellow representing less significant and red more significant. Translation, immune system process and lipid metabolic process were the most enriched biological pathways in stage 2 blood.

Additionally, lipid metabolic process, lipid transport, muscle organ development and cellular amino acid catabolic process were uniquely enriched in stage 2 biological processes.

### Blood Markers for Stage 1 and Stage 2 r-HAT in Malawi

Next, we compared significantly expressed (padj<0.05) CDS in stage 1 (2454 CDS) and stage 2 (850 CDS) blood. We identified 632 CDS that were differentially expressed in both stage 1 and stage 2 r-HAT disease **(Fig 4A)**. Among the 632 CDS, *ZNF354C* was upregulated (log2FC 1.9) in stage 1 only, whereas *TCN1* (log2FC 2.0) and MAGI3 (log2FC 1.4) were upregulated in stage 2 blood only **(Fig 4B)**. Overexpression of *ZNF354C* has a crucial role in inhibition of endothelial cell sprouting (31). *TCN1* expression in blood is negatively associated with poor verbal memory performance (32). Among CDS only expressed in stage 1 blood, 71 genes were upregulated with log2FC >3.0 **(Fig S3A)**. On the other hand, *DMD, NOXRED1, HBB, PROK2, LIMS2, CD14* were the top upregulated (log2FC >1.9) genes significant differentially expressed in stage 2 disease only **(Fig S3B)**. CD14 is an antigen receptor mainly expressed by macrophages during innate immune response and HBB is a crucial for synthesis of ß-globin which form the main structure of the human haemoglobin A (33). To determine the biological processes enriched by genes which were differentially expressed in either stage 1 or stage 2 r-HAT, we also subjected the gene list to ExpressAnalyst in biological pathways. This identified enrichment of circadian rhythm and regulation of translation in stage 2 blood and translation, immune system process, viral process in stage 1 blood among other pathways (**Fig S4A and S4B**).

**Fig 4.**
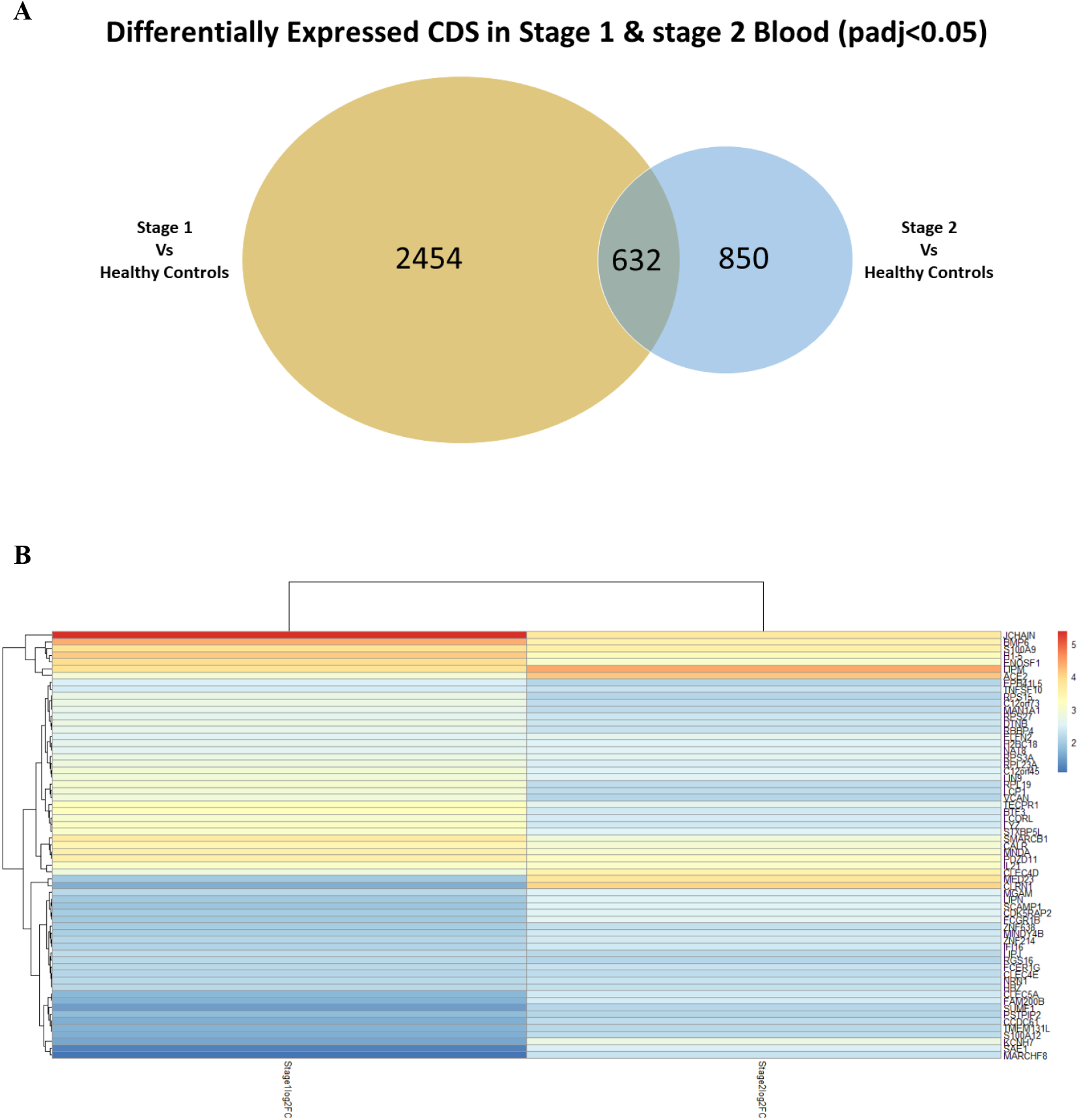
Comparison of protein coding genes differentially expressed in Stage 1 and Stage 2 blood (padj < 0.05). **(A)** Number of DE protein coding genes found in both Stage 1 and Stage 2 cases. **(B)** Hierarchical clustering heatmap of DEGs with log2FC > 2.1 intersecting in Stage 1 and stage 2 blood.

### Neutrophils underlie Differentially Expressed Blood Cells in r-HAT Disease in Malawi

The transcriptional map of human blood cells provides a comprehensive understanding of physiological haematopoiesis (34). We used a custom R script that uses normalised read counts produced by DESeq2 to obtain the proportions of different leukocyte types present in each sample. In a principal component analysis of the data PC1 largely separated cases from controls and explained 25% of the variance in the data (**Fig S5A**). The transformed bulk RNAseq to single cell proportions data had the expected normal distribution (**Fig S5B**). We identified 12 blood cell types with significantly different relative abundance (p<0.05) in r-HAT cases and controls **(Fig 5A, Fig S6 and Table S3)**. Meta-myelocytes (metaN) had the greatest difference in proportions (p<7.4E-6) followed by NKP (p<6.1E-4) and hMDP (p<6.3E-4). Meta-myelocytes are neutrophil precursors and their presence in blood circulation is an indication of severe acute inflammation (35). To understand the immunological pathways involved in r-HAT in Malawi patients, we subjected all upregulated CDS to reactome immune system pathway visualisation (36). This identified neutrophils and macrophages as one of the early responders to trypanosome infection as well (**Fig 5B**).

**Fig 5.**
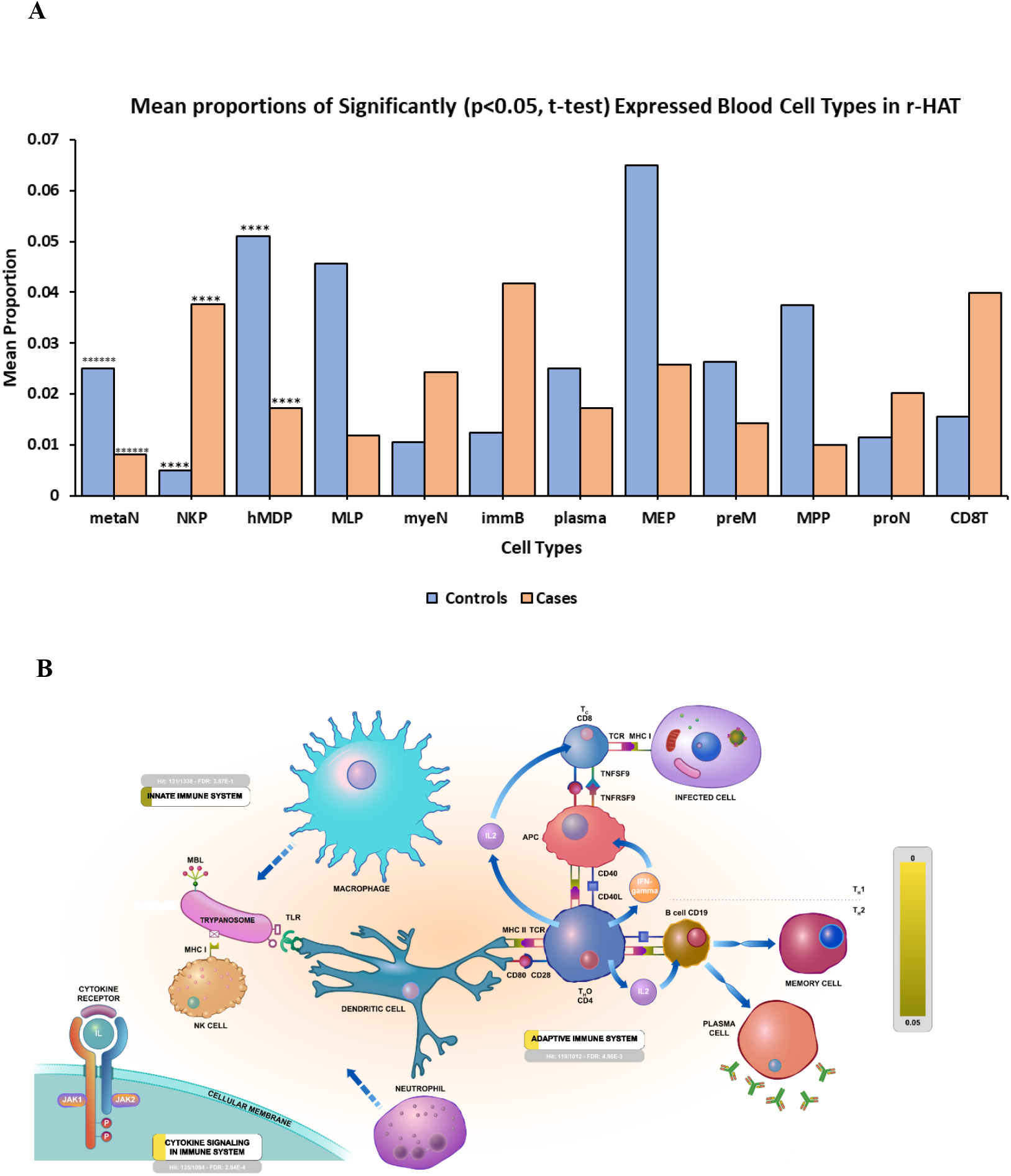
Immune system Blood cells activated in r-HAT. **(A)** Blood cell types that had significantly different proportions in r-HAT cases and controls (p<0.05, t-test). See table S3 for full cell type names. ****p < 6E-04, ******p < 7E-06 **(B)** Visualized output of innate and adaptive immune system pathways interaction in r-HAT cases based on DEGs loaded into Reactome pathway database (36). Yellow represents more significant and darker yellow less significant. Macrophages, neutrophils and NK cells participate in innate immune response which results in significant activation of cytokine signaling pathway (FDR: 2.94E -4). Dendritic cells link innate immune system and activation of adaptive immune system through activation of CD4+ TH cells. Activated CD4 cells release IL2 and IFN-gamma that activates CD8+ T cells and B cells to differentiate into plasma cells for antibody production.

## Discussion

Our previous study showed that clinical presentation of r-HAT in Malawi is focus dependent with most r-HAT cases in Nkhotakota focus presenting with stage 1 disease and in Rumphi focus mostly presenting with stage 2 disease (10).

In this study we have presented transcriptome data from blood of stage 1 and stage 2 r-HAT cases in Nkhotakota and Rumphi foci in Malawi. Nonetheless, blood samples in both stages of r-HAT showed a detectable stratification between cases and healthy controls.

Our data also showed activation of innate immune system in both stage 1 and stage 2 disease. Hematopoietic progenitor neutrophils (Metamyelocytes) were the most significantly (p<7.4E-6) expressed blood cells responding to Tbr infection in humans followed by NKP and hMDP which are all central in coordinating and effecting an innate immune response. Presence of Metamyelocytes in blood is an indication of acute inflammation, which is consistent with proinflammatory profiles in r-HAT (12). Circulating neutrophil life span is about 48hrs, at the same time BMP6 which is involved in cell proliferation was a significant DEG (padj<10E-11) and upregulated in both stage 1 & stage 2 r-HAT. This suggests that innate immune response through neutrophil activation might have a central role in responding to blood parasitaemia in Malawi r-HAT patients. Candidate genes in neutrophil activation have also been identified to respond to *Trypanosoma congolense* infection in cattle (37). Whereas, in mice infected with *T. brucei brucei* (Tbb) neutrophils were recruited at the site of tsetse fly bite but were not able to immobilise motile trypanosomes but aided in the establishment of Tbb blood infection (27). This implicates the dynamic role of neutrophils in responding to various trypanosome parasite infections in different mammalian hosts and future research should consider delineating the role of neutrophils in human Tbr infections.

We also observed upregulation (log2FC 1.9) of CD14 transcripts in blood from stage 2 r-HAT patients. CD14 is involved in activation of macrophages and regulation of macrophage metabolic profiles (38), which was consistent with our finding of activated lipid metabolic process, lipid transport and cellular amino acid metabolic process in stage 2 blood only (**Fig 3B**). Macrophages are involved in clearance of tissue pathogens. At the same time, trypanosomes are known to localise in adipose tissue underneath the skin when there’s an influx of host adaptive immune responses induced by trypanosome variant surface glycoproteins, thereby sustaining host infection in the absence of blood trypanosome parasitaemia (39). Our results might suggest macrophage infiltration in Tbr infected individuals in Malawi, consistent with findings in mice models infected with Tbr (40).

Trypanosome infections are known to disrupt circadian rhythm in vivo and in vitro (23), and here, we found that *CIPC* and *PER1* genes were down regulated and upregulated respectively in stage 1 blood. This suggest that subtle disruption of host circadian system by the trypanosome parasite may start early in infection during hemolymphatic stage, although sleep disturbance is only observed in late stage 2 r-HAT (41, 42).

Comparison of DEGs in stage 1 and stage 2 blood identified *ZNF354C* significant differentially expressed in blood of both stages of r-HAT but upregulated in stage 1 only. Whereas *TCN1* and *MAGI3* were only upregulated in stage 2 blood and neither upregulated nor downregulated in stage 1 blood but significant differentially expressed in both stage 1 and stage 2 compared to healthy controls. These have a diagnostic potential of being used as blood markers to diagnose stage 1 and stage 2 r-HAT cases without need of the invasive lumber puncture collection of CSF, which is currently used for diagnosis of late stage 2 disease. Unlike in a similar study in Uganda r-HAT patients which identified *C1QC, MARCO* and *IGHD3–10* upregulated in both blood and CSF, these transcripts were neither upregulated nor significantly differentially expressed in Malawi r-HAT patients. This supports the need for personalised medicine but not universal medicine in the treatment of r-HAT as infected individuals in different disease focus respond differently to trypanosome infection.

In conclusion, this study has compared transcriptomes differentially expressed and upregulated in blood of stage 1 and stage 2 r-HAT cases in Malawi. We have identified transcripts significant differentially expressed and upregulated in each stage of r-HAT disease. We have identified neutrophils as significant responders of blood trypanosome infection in both stages of the disease, and macrophages as possible responders in patients with late stage disease. We have also identified transcripts that may potentially be used as novel biomarkers in future research for diagnosis of stage 1 and stage 2 r-HAT in Malawi without the need of lumber puncture. Our study has provided insights into human responses to trypanosome infection in Malawi r-HAT patients.

## Methods

### Ethics, Study sites and sample collection

We have recently described r-HAT surveillance and study participants recruitment (10). This study was approved by Malawi National Health Sciences Research Committee (Protocol Number: 19/03/2248). Consent and accent were obtained from each study participant before sample collection. Briefly, sample collection was done during active and passive r-HAT surveillances conducted for 18 months from July 2019 to December 2020. Both r-HAT cases and healthy controls were confirmed to be infected with trypanosome parasites or not by microscopic examination of thick blood films during the surveillance period. Upon obtaining consent, 2ml whole blood samples were collected into Paxgene^®^ tubes from r-HAT cases and matching trypanosome negative healthy individuals and stored at -20ºC until processing. Healthy controls were matched for sex, age group and disease focus. For r-HAT positive individuals, samples were collected before initiation of HAT treatment and all patients were thereafter treated following the national HAT treatment guidelines.

### RNA sequencing and analysis

RNA was extracted from the preserved Paxgene^®^ blood as previously described (43). A minimum of 1µg of total RNA was shipped to the Center for Genomics Research at the University of Liverpool for sequencing. Samples were checked for quality using an Agilent Bioanalyzer and samples with RNA < 1µg were excluded. Libraries were prepared from total RNA using the QIASeq FastSelect rRNA, Globin mRNA depletion and <u>NEBNext Ultra II Directional RNA Library Prep Kit</u> and were sequenced to a target depth of 30 million reads on the Illumina® NovaSeq (100 million reads for samples infected with Tbr parasites). FASTQ reads were aligned to the GRCh38 release 84 human genome sequence obtained from Ensembl (44) using HiSat2 (45) and annotated using the *Homo sapiens* GRCh38.104.gtf file from Ensembl. Genes that were differentially expressed between phenotypes were identified using DEseq2 (46). The proportions of different cell types in each sample were estimated using Bisque (47). Single cell reference sequence data from bone marrow and peripheral blood from Chinese donors was obtained from 7551 individual human blood cells representing 32 immunophenotypic cell types (34). Network analysis of enriched genes was done using XGR (48), InnateDB (49) and ExpressAnalyst (28).

## Author Contributions

**Peter Nambala:** Conceptualization, Methodology, Investigation, Formal analysis, Writing - original draft. **Harry Noyes:** Conceptualization, Methodology, Formal analysis, Writing - review & editing. **Julius Mulindwa:** Conceptualization, Writing - review & editing, Methodology, Formal analysis, Supervision. **Joyce Namulondo:** Formal analysis. **Oscar Nyangiri:** Formal analysis. **Enock Matovu:** Conceptualization, Supervision. **Annette MacLeod:** Conceptualization. **Janelisa Musaya:** Conceptualization, Writing - review & editing, Methodology, Supervision, Formal analysis.

## Acknowledgement

We would like to acknowledge Nkhotakota and Rumphi district health offices for their assistance in sample collection.

## Supplementary Tables

**Table S1.**
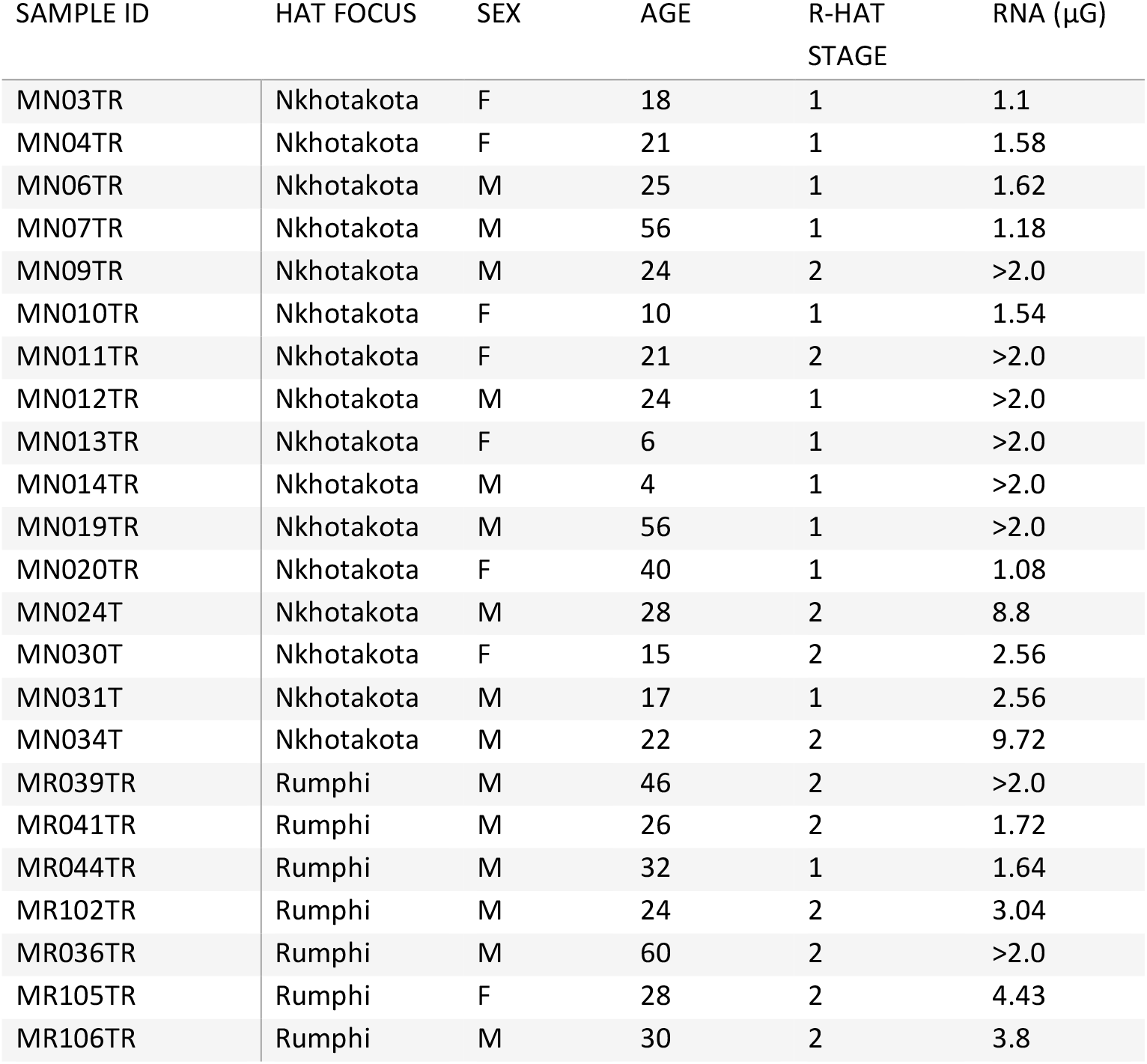
Summary on blood samples from r-HAT cases used for RNA sequencing.

| SAMPLE ID | HAT FOCUS | SEX | AGE | R-HAT<br>STAGE | RNA (μG) |
| --- | --- | --- | --- | --- | --- |
| MN03TR | Nkhotakota | F | 18 | 1 | 1.1 |
| MN04TR | Nkhotakota | F | 21 | 1 | 1.58 |
| MN06TR | Nkhotakota | M | 25 | 1 | 1.62 |
| MN07TR | Nkhotakota | M | 56 | 1 | 1.18 |
| MN09TR | Nkhotakota | M | 24 | 2 | >2.0 |
| MN010TR | Nkhotakota | F | 10 | 1 | 1.54 |
| MN011TR | Nkhotakota | F | 21 | 2 | >2.0 |
| MN012TR | Nkhotakota | M | 24 | 1 | >2.0 |
| MN013TR | Nkhotakota | F | 6 | 1 | >2.0 |
| MN014TR | Nkhotakota | M | 4 | 1 | >2.0 |
| MN019TR | Nkhotakota | M | 56 | 1 | >2.0 |
| MN020TR | Nkhotakota | F | 40 | 1 | 1.08 |
| MN024T | Nkhotakota | M | 28 | 2 | 8.8 |
| MN030T | Nkhotakota | F | 15 | 2 | 2.56 |
| MN031T | Nkhotakota | M | 17 | 1 | 2.56 |
| MN034T | Nkhotakota | M | 22 | 2 | 9.72 |
| MR039TR | Rumphi | M | 46 | 2 | >2.0 |
| MR041TR | Rumphi | M | 26 | 2 | 1.72 |
| MR044TR | Rumphi | M | 32 | 1 | 1.64 |
| MR102TR | Rumphi | M | 24 | 2 | 3.04 |
| MR036TR | Rumphi | M | 60 | 2 | >2.0 |
| MR105TR | Rumphi | F | 28 | 2 | 4.43 |
| MR106TR | Rumphi | M | 30 | 2 | 3.8 |

**Table S2:**
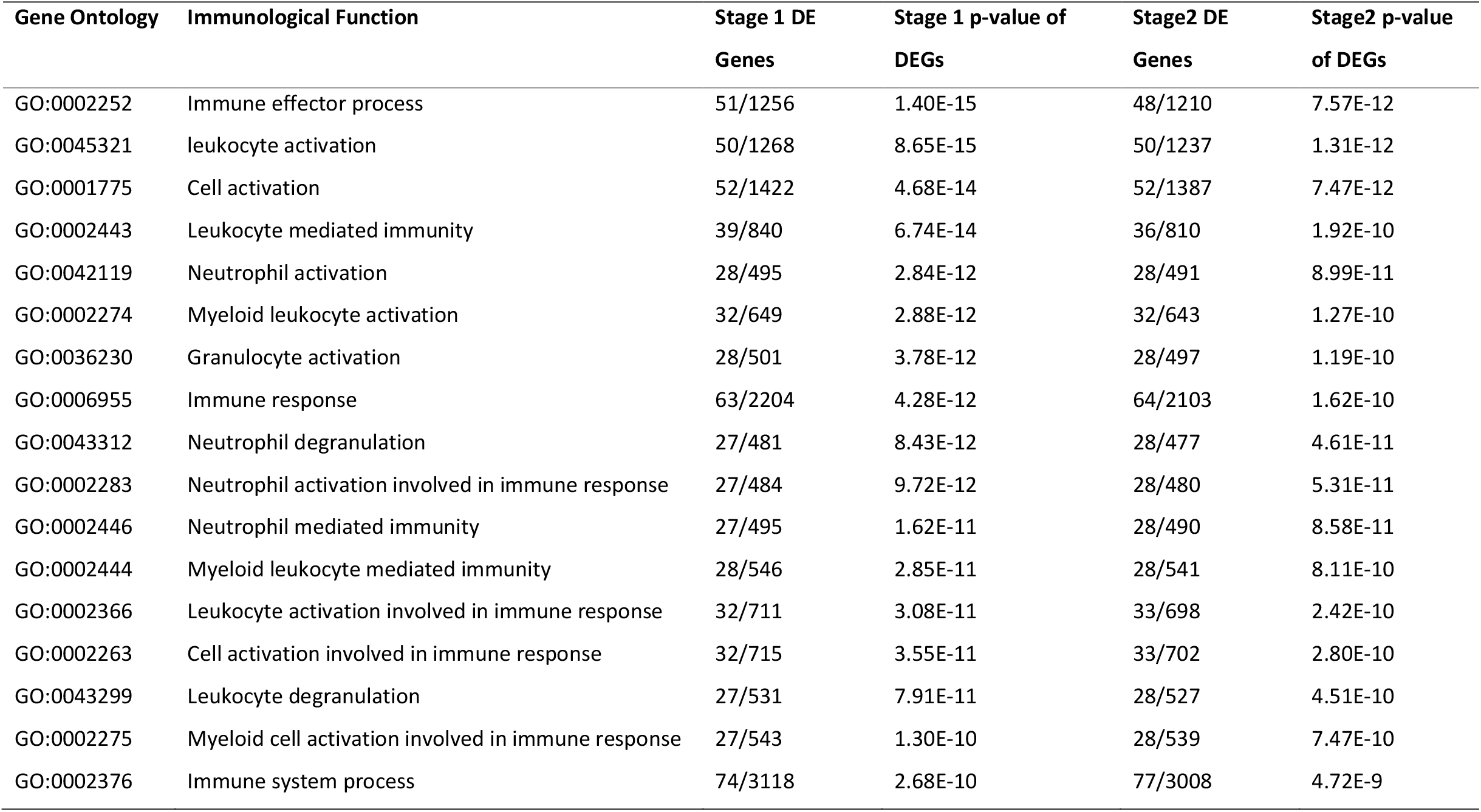
PCA2GO Functional Annotation of Immune biological functions significantly enriched p<10E-10) in Stage 1 and stage 2 r-HAT cases. DEGs = Differentially expressed genes.

**Table S3:** Mean proportion by and p-values (t-test) of cell type for cases and controls.

| Cell Type<br>Abbreviation | Full Cell Type Name | Mean of<br>Cases | Mean of<br>Controls | Mean<br>Difference | p-Value (t-test<br>of CAS vs CON) |
| --- | --- | --- | --- | --- | --- |
| metaN | Meta-myelocyte | 0.008144 | 0.024921 | 0.016776642 | 0.000007356 |
| NKP | Natural killer progenitor | 0.0376 | 0.005025 | 0.032574543 | 0.000608379 |
| hMDP | Human monocyte-dendritic<br>progenitor | 0.017143 | 0.050956 | 0.033812759 | 0.000625394 |
| MLP | Multi-lymphoid progenitor | 0.01181 | 0.045602 | 0.033791638 | 0.001753953 |
| myeN | Myelocyte | 0.024252 | 0.010507 | 0.013745726 | 0.004156435 |
| immB | Immature B lymphocyte | 0.04164 | 0.012412 | 0.029227973 | 0.006327324 |
| plasma | Plasma cells | 0.017188 | 0.025015 | 0.007826678 | 0.010696056 |
| MEP | Megakaryocyte–erythroid progenitor | 0.025675 | 0.064903 | 0.039228534 | 0.011838218 |
| preM | Pre-monocyte | 0.014212 | 0.026374 | 0.012161921 | 0.013904749 |
| MPP | Multi-potent progenitor | 0.009874 | 0.037404 | 0.027530134 | 0.014930755 |
| proN | Pro-myelocyte | 0.02025 | 0.011471 | 0.008779258 | 0.029865929 |
| CD8T | CD8 T-helper | 0.039784 | 0.015558 | 0.024226424 | 0.031372518 |
| regB | regulatory B cells | 0.017841 | 0.046317 | 0.028475456 | 0.05464077 |
| kineNK | Cytokine natural killer | 0.019987 | 0.029325 | 0.009337392 | 0.056628973 |
| matureN | Mature neutrophils | 0.008926 | 0.003806 | 0.005120196 | 0.06528435 |
| GMP | granulocyte/monocyte progenitor | 0.011928 | 0.000471 | 0.011456411 | 0.070773567 |
| CD4T | CD4 T-helper | 0.047214 | 0.022466 | 0.02474757 | 0.076720819 |
| CMP | Multipotent common myeloid<br>progenitor | 0.061713 | 0.027791 | 0.033921159 | 0.082486356 |
| ery | Erythrocytes | 0.277417 | 0.226845 | 0.050572677 | 0.105307824 |
| preB | Precursor B lymphocyte | 0.017942 | 0.032148 | 0.014205465 | 0.159322202 |
| claM | Classical monocytes | 0.018357 | 0.012335 | 0.006021536 | 0.215157411 |
| nonM | Non-classical monocytes | 0.020461 | 0.027314 | 0.006852195 | 0.224520598 |
| toxiNK | Cytotoxic natural killer | 0.004444 | 0.006744 | 0.002299807 | 0.238366919 |
| interM | Intermediate monocyte | 0.016918 | 0.010066 | 0.006852145 | 0.24649224 |
| LMPP | Lympho-myeloid primed progenitor | 0.04841 | 0.073465 | 0.025054531 | 0.339792864 |
| memB | Memory B lymphocytes | 0.030201 | 0.021041 | 0.009160112 | 0.404516409 |
| HSC | Hematopoietic stem cells | 0.01893 | 0.013814 | 0.005115975 | 0.490063317 |
| cMOP | Common monocyte progenitor | 0.016631 | 0.012838 | 0.003792378 | 0.495129766 |
| CLP | Common lymphoid progenitor | 0.011293 | 0.008161 | 0.003131858 | 0.584340877 |
| proB | Progenitor B lymphocytes | 0.020959 | 0.025184 | 0.00422504 | 0.584554591 |
| naïB | Naïve B lymphocytes | 0.039013 | 0.043473 | 0.004459464 | 0.686283191 |
| BNK | Peripheral Blood natural killer | 0.023842 | 0.02625 | 0.002408283 | 0.846255276 |

## Supplementary Figures

**Fig S1.**
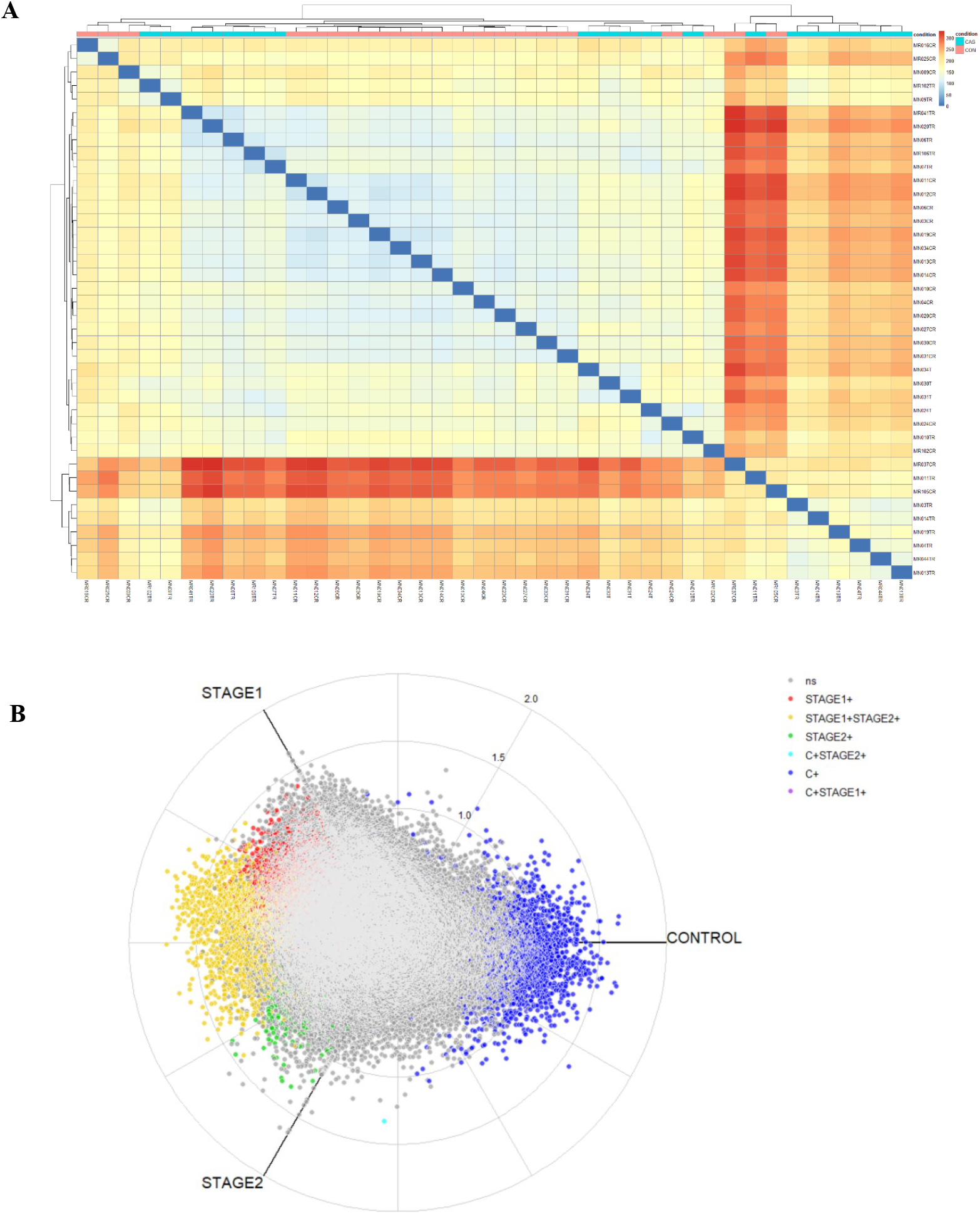
Differential gene expression in r-HAT cases vs controls. **(A)** Sample to sample hierarchical clustering heatmap with complete linkage of cases vs controls. **(B)** Radial plot of the distribution and interception of DEGs in Stage1 and Stage 2 blood vs control blood. Grey color represents genes that were not significant; red represents genes enriched in stage 1 only; green represents genes enriched in stage 2 only; blue represents genes enriched in controls only; light blue genes in cases and control; pink represents genes enriched in both stage1 and controls; and yellow represents genes enriched in both stage 1 and stage 2 blood

**Fig S2.**
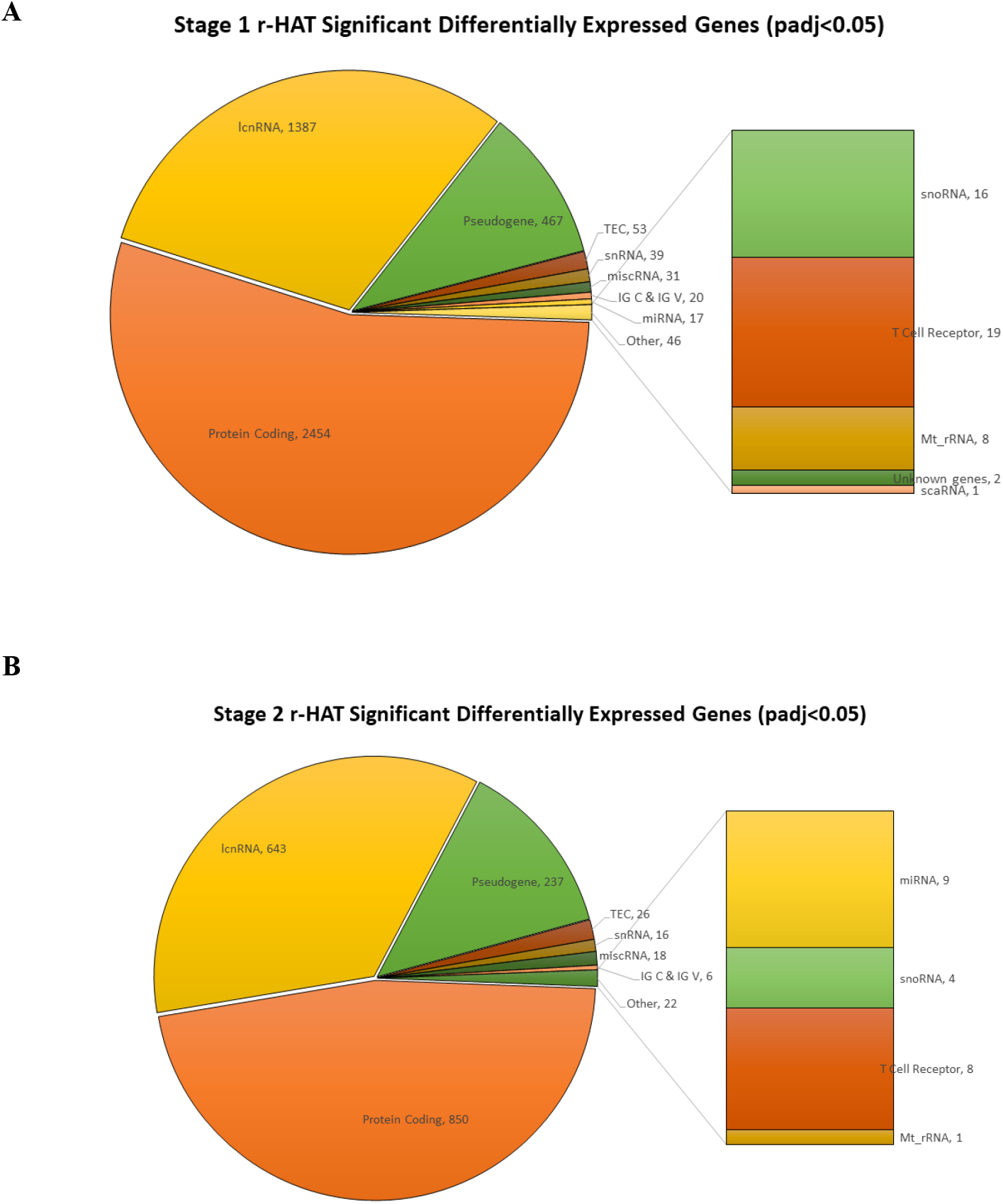
Significant differentially expressed genes. Gene types that were significant differentially expressed in Stage 1 **(A)** and Stage 2 **(B)** r-HAT. Protein coding genes were the most differential expressed followed by lcnRNA and pseudogenes in both stage 1 and 2 r-HAT.

**Fig S3.**
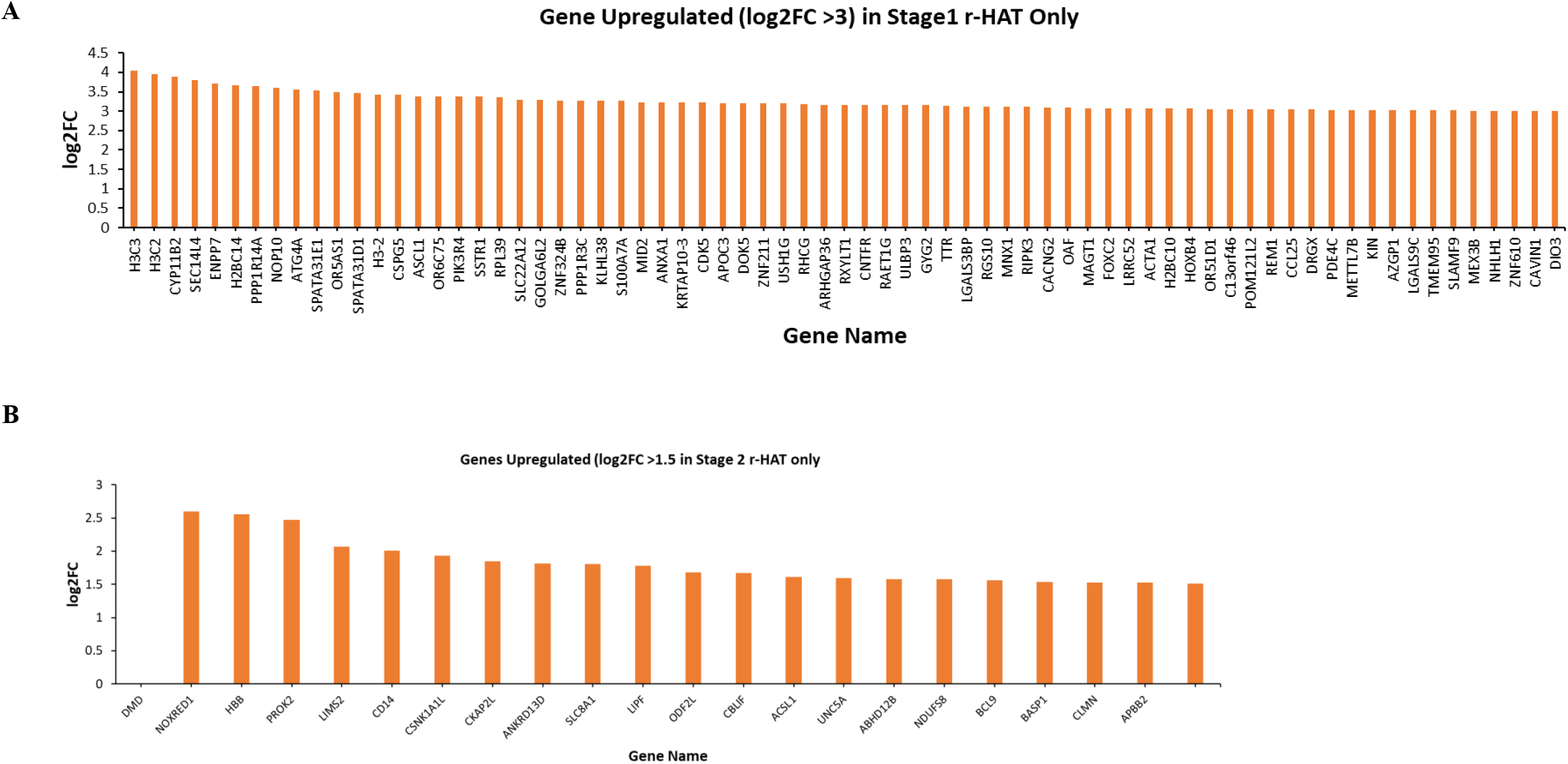
3 List of genes specifically upregulated in blood of stage 1 (log2FC > 3.0) only **(A)**, and **(B)** in stage 2 (B, log2FC > 1.5) only.

**Fig S4.**
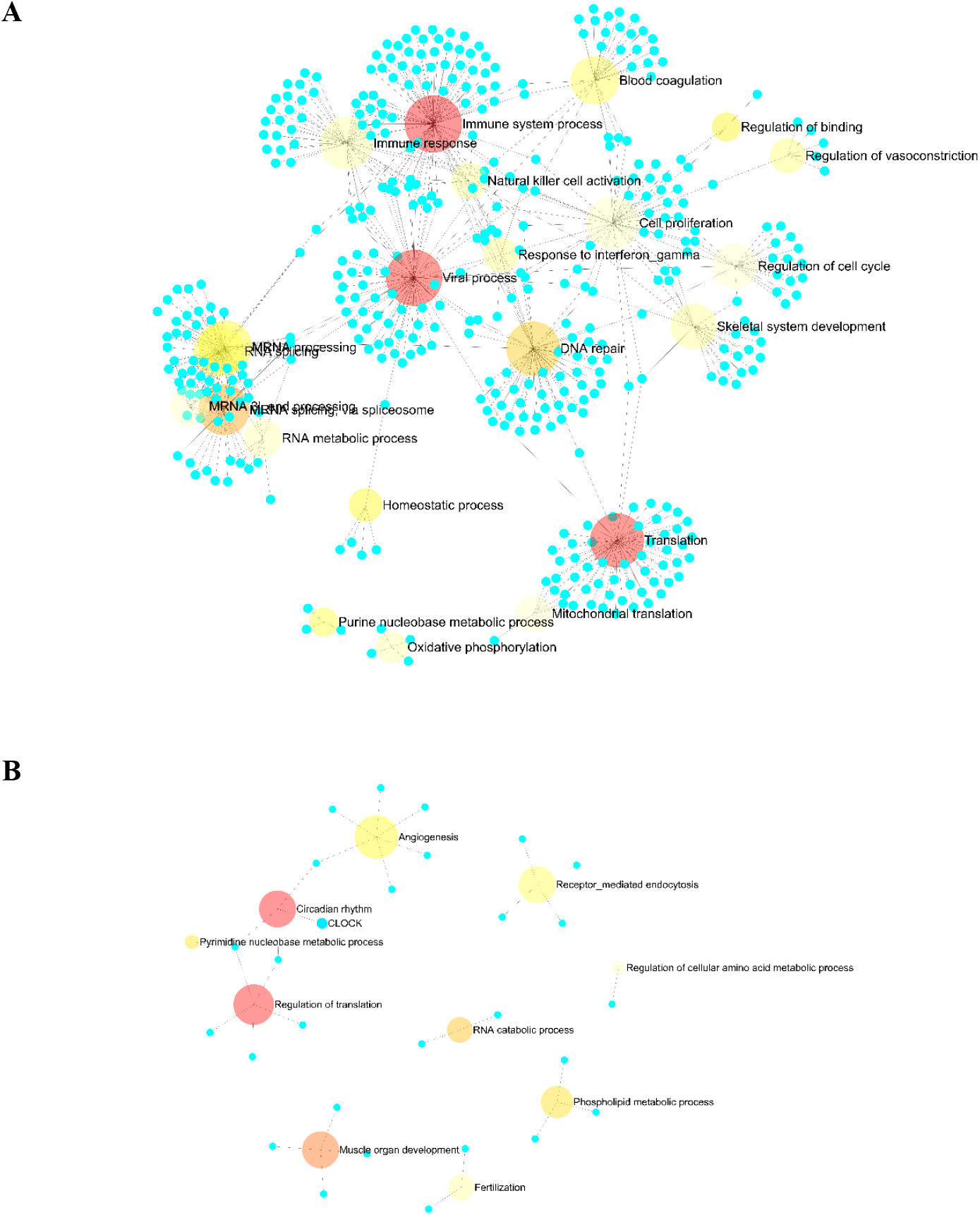
Biological pathways enriched in genes DE in Stage 1 blood only **(A)** and in Stage 2 blood only **(B)**. Images generated by ExpressAnalyst.

**Fig S5.**
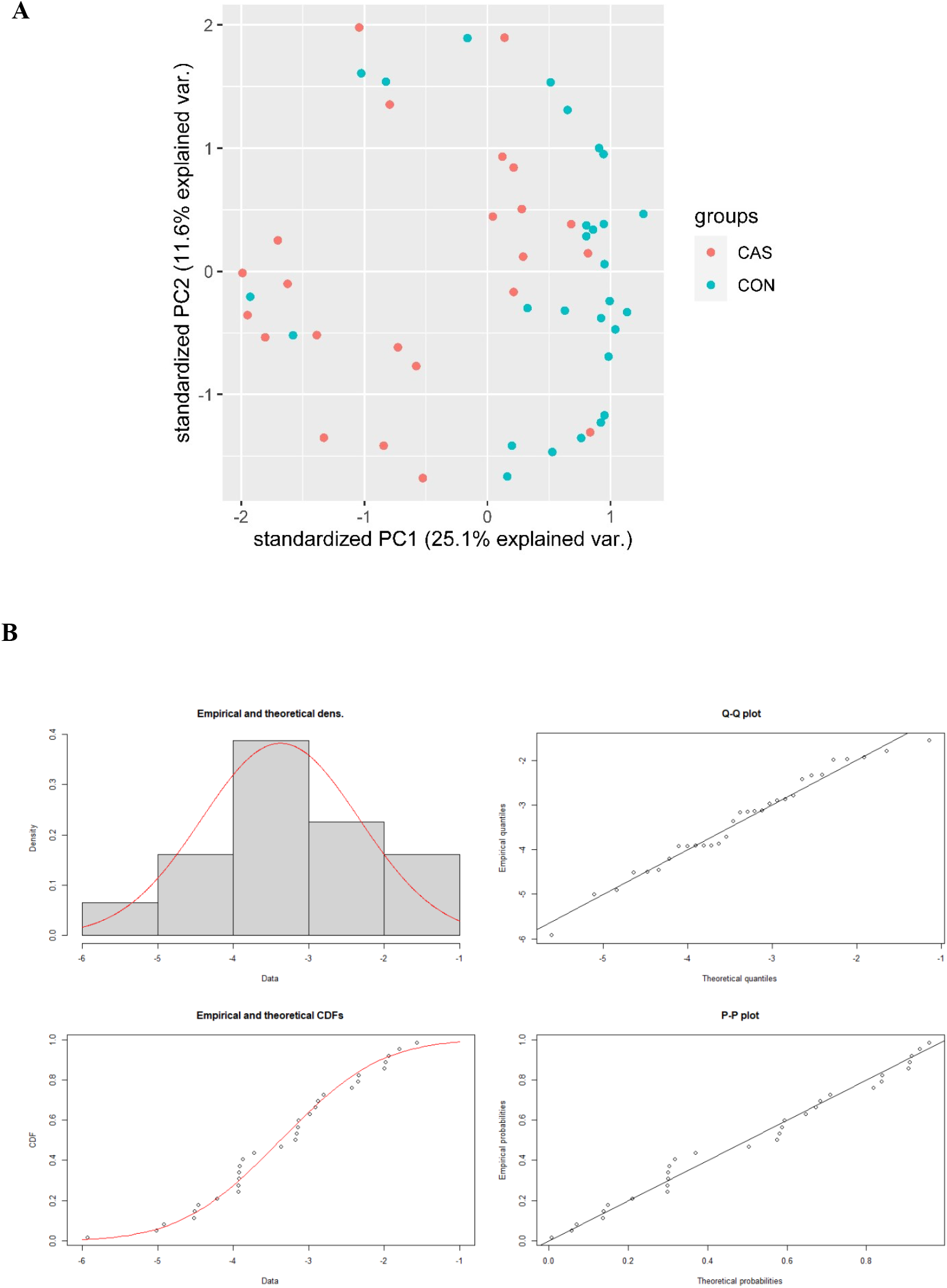
Stratification of single blood cells in cases versus controls. **(A)** 1^st^ and 2^nd^ principal components of the proportions of different cell types by phenotype. The cases and controls are separated along PC1. **(B)** Plots showing normal distribution of the transformed bulk RNA-Seq data into blood single cell RNA-seq data.

**Fig S6.**
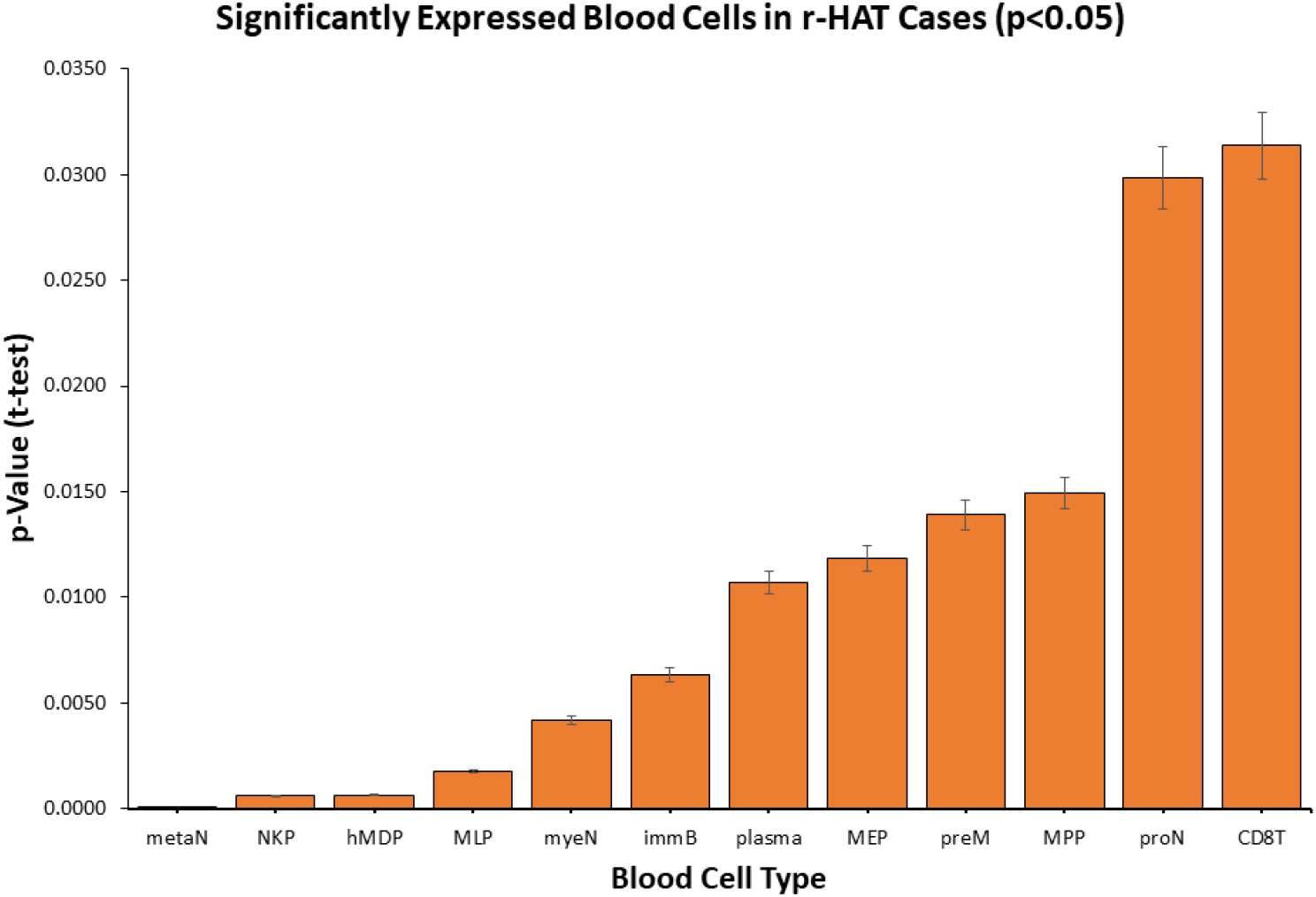
Blood cell types that had significantly different proportions in r-HAT cases and controls (p<0.05, t-test). See table S3 for full cell type names.

